# Frequent translation of small open reading frames in evolutionary conserved lncRNA regions

**DOI:** 10.1101/348326

**Authors:** Jorge Ruiz-Orera, M.Mar Albà

## Abstract

The mammalian transcriptome includes thousands of transcripts that do not correspond to annotated protein-coding genes. Although many of these transcripts show homology between human and mouse, only a small proportion of them have been functionally characterized. Here we use ribosome profiling data to identify translated open reading frames, as well as non-ribosomal protein-RNA interactions, in evolutionary conserved and non-conserved transcripts. We find that conserved regions are subject to significant evolutionary constraints and are enriched in translated open reading frames, as well as non-ribosomal protein-RNA interaction signatures, when compared to non-conserved regions. Translated ORFs can be divided in two classes, those encoding functional micropeptides and those that show no evidence of protein functionality. This study underscores the importance of combining evolutionary and biochemical measurements to advance in a more complete understanding of the transcriptome.

## INTRODUCTION

The advent of high-throughput genomic technologies has revealed that mammalian transcriptomes are more complex than initially thought (Carninci et al., 2005; Kapranov et al., 2007; Okazaki et al., 2002; Ponjavic et al., 2007). One of the most intriguing findings has been the discovery of thousands of expressed loci that lack conserved or long ORFs (Cabili et al., 2011; Liu et al., 2012; Okazaki et al., 2002; Pauli et al., 2012; Ponting et al., 2009; Ulitsky and Bartel, 2013). These transcripts, commonly denominated long non-coding RNAs (lncRNAs), share many of the features of coding mRNAs, such as the presence of a polyadenylated tail and a multi-exonic structure (Consortium et al., 2007). Some lncRNAs may have originated as a result of bidirectional transcription from promoters (Lepoivre et al., 2013) or enhancers (Hon et al., 2017; Li et al., 2016), whereas others may have been born thanks to the fortuitous appearance of weak promoters in the genome (Ruiz-Orera et al., 2015).

The function of lncRNAs is a matter of intense debate. In general, lncRNAs display high evolutionary turnover (Kutter et al., 2012; Neme and Tautz, 2016), and show very weak sequence constraints according to single nucleotide polymorphism data (Wiberg et al., 2015). This is consistent with the idea that many lncRNAs are not functional but a result of the high transcriptional activity of the genome (Brosius, 2005; Struhl, 2007; Wang et al., 2004). However, some lncRNAs have been shown to regulate gene expression through interactions with specific proteins in the nucleus or the cytoplasm (Gong et al., 2012; Han et al., 2010; Ribeiro et al., 2017) or with the chromatin (Ponting et al., 2009), even if expressed at very low levels (Seiler et al., 2017). Currently, the fraction of lncRNAs that are functional is unknown.

Not surprisingly, functional lncRNAs often contain short conserved sequence segments that are required for their function (Kapusta and Feschotte, 2014; Ulitsky, 2016). Although very few lncRNAs display deep conservation in vertebrates, hundreds of lncRNAs show conservation between human and mouse (Necsulea et al., 2014; Ulitsky et al., 2011). The conserved sequence patches tend to be short and 5’-biased (Hezroni et al., 2015). According to polymorphism data, the evolution of conserved lncRNAs tends to be more constrained than the evolution of non-con-served lncRNAs (Wiberg et al., 2015), indicating that the former lncRNAs are enriched in functional sequences.

Ribosome profiling sequencing data (Ribo-Seq), which captures RNA-ribosome interactions but also other types of RNA-protein interactions (Ingolia et al., 2009; Ji et al., 2016), offers new opportunities to investigate the properties of conserved *versus* non-conserved lncRNA regions. Remarkably, Ribo-Seq has revealed the existence of thousands of translated open reading frames (ORFs) in lncRNAs (Bazzini et al., 2014; Castaneda et al., 2014; Ingolia et al., 2011; Ji et al., 2015; Ruiz-Orera et al., 2014). Some of them correspond to small functional proteins which have been missed by gene annotation pipelines, such as myoregulin (Mrln) (Anderson et al., 2015) or TUNAR (Megamind) (Lin et al., 2014). Other ORFs are likely to translate non-functional peptides according to polymorphism data (Ruiz-Orera et al., 2014, 2018). The footprints of non-ribosomal ribonucleoprotein particles have also been detected on some functional lncRNAs because of the distinctive length of the associated Ribo-Seq reads (Ingolia et al., 2014).

Here we investigate the presence of Ribo-Seq-related signatures, as well as other annotated features such as putative promoter sequences, in lncRNAs sequences that are conserved between mouse and human. We find that conserved regions contain an excess of promoter sequences, translated ORFs and non-ribosomal ribonucleoprotein particles when compared to non-conserved regions.

## RESULTS

### Conserved regions in lncRNAs are enriched in translation and regulatory signatures

We searched for matches of the complete set of Ensembl mouse annotated transcripts against the human transcriptome using BLASTN (E-value < 10^−5^) (Altschul et al., 1997). The human transcriptome was obtained using high coverage RNA sequencing data (RNA-Seq) from different tissues (Ruiz-Orera et al., 2015). We detected 19,779 conserved protein-coding genes (codRNAs) and 1,547 conserved lncRNAs, containing at least one conserved region in humans. The conserved regions were in general shorter in lncRNAs than in codRNAs (median length of 163 and 343 nucleotides, respectively; Additional file 1: Figure S1 for more details).

Next, we focused on genes expressed that were significantly expressed in brain tissue, using a high coverage ribosome profiling dataset from mouse hippocampus (Cho et al., 2015). We detected significant expression for 13,081 conserved codRNAs and 289 conserved lncRNAs, including 444 conserved lncRNA regions (Additional file 1: Table S1). The set of conserved lncRNAs was enriched in functionally characterized lncRNAs (27 cases, see Additional File 2); the only exceptions were *Firre*, *Adapt33*, and *Snhg6*. The percentage of genes with at least one conserved region was 98% for codRNAs and 40.88% for lncRNAs (Figure 1a). In terms of total length, 8.50% of the total lncRNA sequence (25.39% in the case of functionally characterized lncRNAs), and 61.62% of the codRNA sequence, was conserved (Figure 1a).

**Figure 1.**
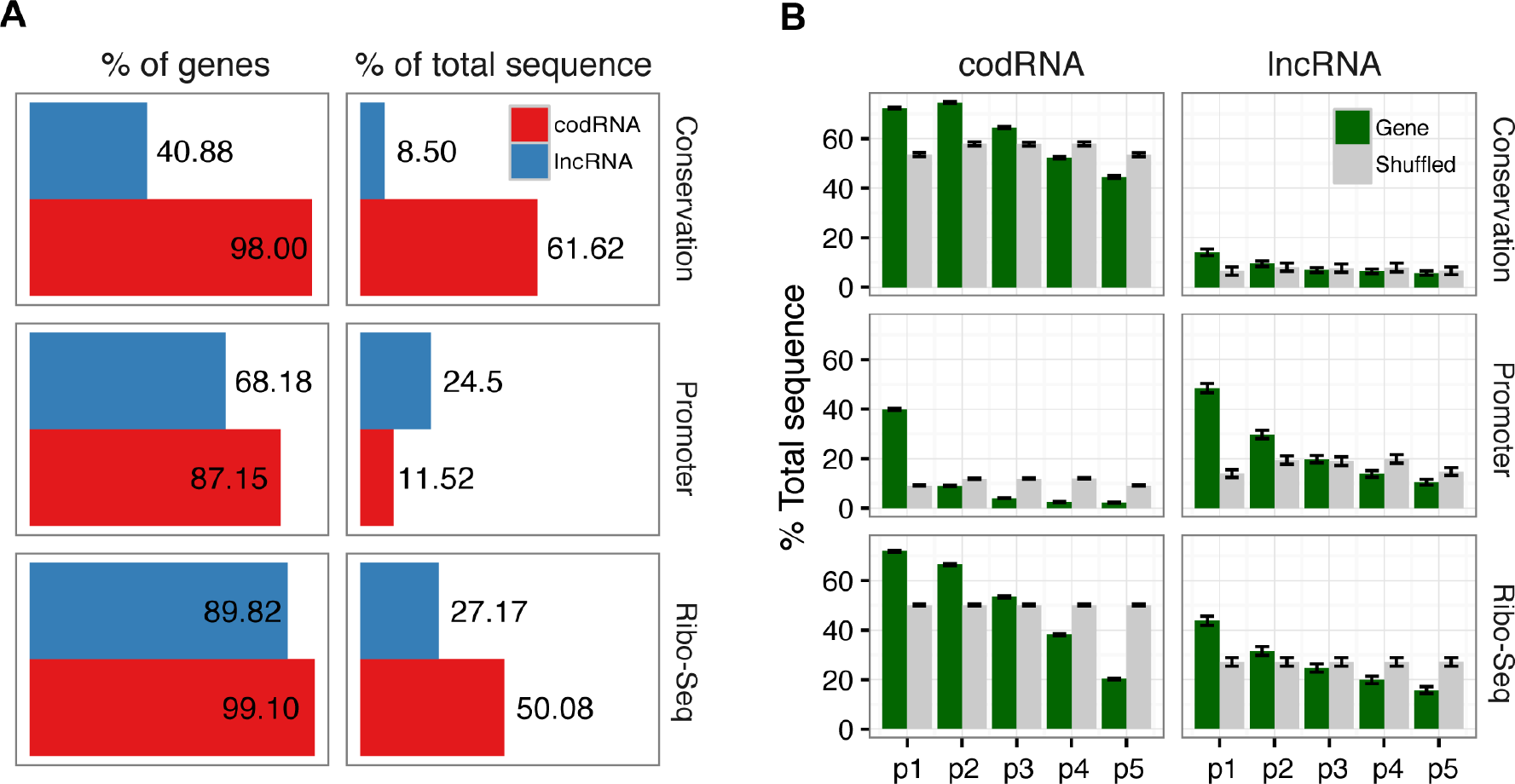
Transcriptome-wide identification of conserved sequences, promoters, and Ribo-Seq associations. **A.** Fraction of mouse genes that showed conservation in human using BLASTN (Conservation), that overlapped with annotated promoter regions (Promoter), or that were covered by Ribo-Seq reads (Ribo-Seq). The percentage of genes with at least one feature, and the total sequence covered, are indicated. Data is for expressed codRNAs and lncRNAs in hippocampus (sequences with a minimum RNA-Seq coverage of 56.38 reads/kb). **B.** Analysis of feature coverage in equally-sized fractions of the genes, from 5’ (p1) to 3’ (p5). Grey bars represent the mean proportion of a shuffled control where the different features per gene were randomly shuffled along the sequence 1000 times. Error bars represent the standard error of the proportion.

We observed that the transcripts frequently overlapped sequences annotated as promoters by Ensembl (Zerbino et al., 2015). This affected 87.15% of codRNAs and 68.18% of lncRNAs. In relative terms, lncRNAs were more extensively covered by promoters (24.50% of total sequence) than codRNAs (11.52% of total sequence) (Figure 1a), and promoter regions were biased towards the 5′ end of lncRNAs (Figure 1b). The relatively high overlap of promoters with lncRNAs could be explained by their short size compared to codRNAs; when we focused on the 5′-most 200 nucleotides of the transcript the percentage of sequence covered by promoters was actually higher for codRNAs than for lncRNAs (80.46% *versus* 63.38%). We observed a strong enrichment of promoters in conserved regions: promoters occupied nearly 54% of the total lncRNA conserved sequence, compared to only ~22% of the non-conserved one (Figure 2b).

**Figure 2.**
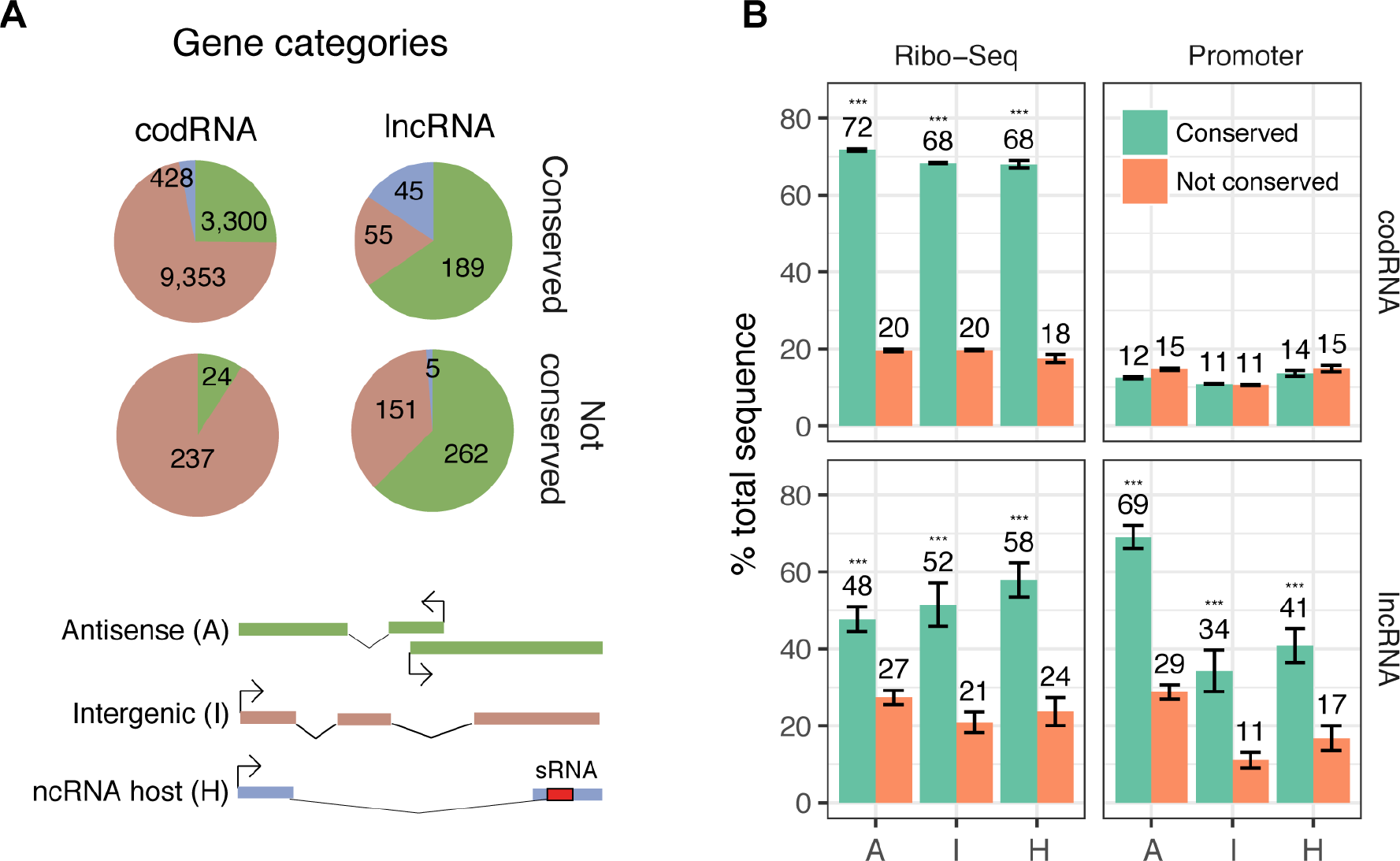
Effect of conservation across lncRNA types. **A.** Number and fraction of different categories based on position and sequence features. Antisense: Exonic overlap, expression on a bidirectional promoter, and/or annotated as antisense; ncRNA host: Genes with at least one found small RNA sequence in the exonic region; intergenic: rest of genes. Conserved genes are enriched in antisense and ncRNA host genes. **B.** Percentage of total sequence that is covered by Ribo-Seq reads (1 or more reads), and annotated promoter cores, for conserved and non-conserved regions in codRNAs and lncRNAs. Conserved lncRNA regions showed a significantly higher proportion of all features compared to not conserved regions or expected randomly (Test of equal proportions; * p-value < 0.05; *** p-value < 10^−5^). Error bars represent the standard error of the proportion. Categories: A: Antisense; I: Intergenic; H: ncRNA host.

Next, we investigated the presence of ribosome profiling (Ribo-Seq) signals on the transcripts. We observed that most codRNAs (99.10%) and lncRNAs (89.92%) were covered by at least 1 Ribo-Seq read. When looking at the total sequence length, 50.08% of the codRNA sequence and 27.17% of the lncRNA sequence was covered by Ribo-Seq reads (Figure 1a). These results are in line with recent reports of a relatively high coverage of lncRNAs by Ribo-Seq reads (Ingolia et al., 2011; Ruiz-Orera et al., 2014). The Ribo-Seq reads showed a clear 5′ bias, both in conserved and non-conserved regions (Figure 1b, Additional file 1: Figure S2).

To account for the fact that some regions might be conserved because of antisense overlaps with protein-coding exons, we did the same analysis without considering any overlapping region, reaching similar conclusions of higher density of Ribo-Seq reads in conserved *versus* nonconserved regions (Additional file 1: Figure S3). Moreover, even though conserved regions had higher RNA-seq and Ribo-Seq read coverage than non-conserved ones (Additional file 1: Figure S4), the same effect could be observed for different expression level intervals, indicating that the trend was robust to variations in the amount of the transcript (Additional file 1: Figure S5). Ribo-Seq data from human and rat brain tissues, for the corresponding genomic syntenic sequences, yielded very similar results (Additional file 1: Figure S6).

### Consistent results across lncRNA types

We divided lncRNAs come in two groups, intergenic lncRNAs (I, Figure 2a) and antisense lncRNAs (A, figure 2a). Intergenic lncRNAs were completely independently loci. Antisense lncRNAs included those lncRNAs annotated as antisense in Ensembl, as well as any other lncRNA whose transcription start site was located less than 2Kb from the TSS of another gene in antisense orientation and/or had antisense exonic overlap with another gene. We also found 50 annotated lncRNAs with embedded short non-coding RNAs in the exons (they contain 33 miRNAs, 41 snoRNAs, and 32 miscRNAs); we termed this class ncRNA host (H, Figure 2a).

Many of these lncRNAs are known to be processed to form small conserved RNA molecules, as it occurs with the 3’ tail of *Malat* (Wilusz et al., 2008), although in other cases (f.e. *Slert*) the presence of the sRNA-like sequence in the lncRNA enables the biogenesis and translocation of the transcript (Xing et al., 2017). For comparative purposes, we classified the genes annotated as coding in the same three categories as the lncRNAs. We observed that the relative frequency of antisense genes was much higher in lncRNAs than in codRNAs (451 out of 707 *versus* 3,324 out of 13,342).

The class defined as ncRNA host was strongly enriched in conserved sequences when compared to the other two lncRNA classes (Figure 2a). Overall, 90% of the mouse ncRNA host sequences showed significant conservation in the human transcriptome, whereas this fraction was 42% for antisense lncRNAs and 27% for intergenic lncRNAs. Promoter sequences and Ribo-Seq mappings were more abundant in conserved than in non-conserved regions for all three lncRNA classes (Figure 2b). The main differences between the classes were an excess of promoter sequences in antisense lncRNAs and increased Ribo-Seq signal in conserved ncRNA host. When the three classes of lncRNAs were taken together, the fraction of regions covered by Ribo-Seq reads was about double for conserved than for non-conserved regions (51.7% versus 24.9%, Test of equal proportions, p-value < 10^−5^).

### Conserved lncRNA sequences are under selection

Although mouse and human are relatively distant species (~ 90 Million years) (Hedges et al., 2015), some sequence segments may be conserved purely by chance. In order to estimate the expected degree of conservation between mouse and human transcripts in the absence of selection we run sequence evolution simulations using Rose (Stoye et al., 1998). In particular, we simulated the evolution of lncRNAs along the mouse and human branches under no evolutionary constraints. Subsequently we performed BLASTN searches of the evolved mouse sequences against the set of evolved human sequences (see Methods for more details). We could find BLASTN homology hits for about 56.2% of the evolved lncRNA sequences. The fact that this fraction is larger than the observed one for real lncRNAs (40.9%, Figure 1a) supports the idea that a fraction of the current mouse lncRNAs have originated after the split with the human lineage.

Next, we used the sequence alignments obtained with BLASTN to estimate the number of substitutions per site (*k*) using the PAML package (Yang, 2007), in different sequence sets. In alignments of size equal or longer than 300 nucleotides, the computed *k* was similar to the expected 0.51 substitutions per site for regions evolving under no constraints (See Methods for more details). Using the same length cut-off the computed *k* for real lncRNAs was 0.25 and hence significantly lower than the expected under no constraints (Wilcoxon test, p-value < 10^−5^). This indicates that purifying selection is acting on lncRNAs containing regions that are conserved between mouse and human.

Alignments shorter than 300 nucleotides tended to give estimates of *k* lower than 0.51 even in the case of the simulated sequences, which was not initially expected. In this size range we observed a positive relationship of *k* with alignment length, with shorter sequences showing lower *k* (Additional file 1: Figure S7). This indicated that short sequenced needed to have a higher percent identity to be detected as significant by BLAST. As many conserved sequences in lncRNAs were lower than 300 nucleotides (Additional file 1: Figure S1) we modeled the effect of length on *k* using two different log-linear regression models, one for short (< 300 nt) and one for long (≥ 300 nt) sequences, using the data from the sequence evolution simulations. This allowed to predict an expected *k* (*k*_<u>*e*</u>_) given a sequence alignment length, which we used to normalized the observed *k* for real sequences (*k*_*o*_/*k*_*e*_).

The *k*_*o*_/*k*_*e*_ was significantly lower in all three categories of lncRNAs than in the neutrally evolved sequences (Figure 3, Wilcoxon test, p-value < 10^−5^), consistent with selection acting on at least some of the lncRNAs. The intensity of the selection signal, as measured with the *k*_*o*_/*k*_*e*_ ratio, was similar for conserved segments in functionally characterized and uncharacterized lncRNAs (median 0.49 and 0.50, respectively). Coding sequences and ncRNA host transcripts also showed clear selection signals (median *k*_*o*_/*k*_*e*_ 0.37 and 0.46, respectively). We also calculated separated values for regions corresponding to Ensembl annotated promoter regions, which spanned 34-69% of the conserved lncRNA regions (Figure 2b). Although conserved promoter regions had a somewhat lower *k*_*o*_/*k*_*e*_ than the rest of conserved lncRNA sequences (median 0.47 *versus* 0.58), the signal of purifying selection continued to be very clear after eliminating the promoters.

**Figure 3.**
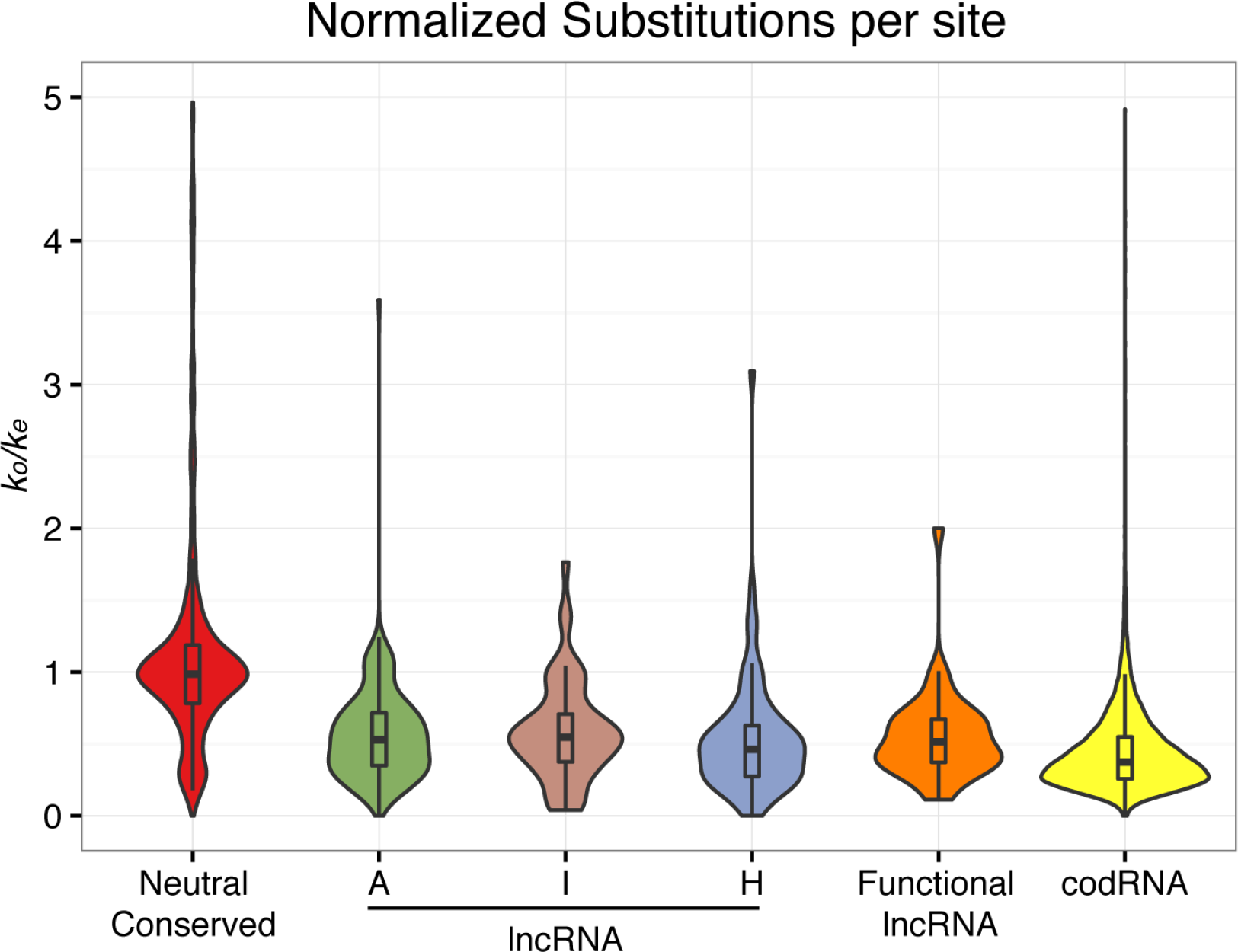
Distribution of normalized substitution rates (*k*_*o*_/*k*_*e*_) between human and mouse sequences with BLAST-based homology. The number of substitutions per site was estimated (*k*_*o*_) in the regions with BLAST hits with baseml under the REV nucleotide substitution model and normalized by dividing it by the expected *k* under neutrality for different length intervals (*k*_*e*_). Neutral conserved: simulated neutrally evolving sequences with BLASTN matches (2,736 regions); lncRNA: lncRNAs with BLASTN matches; A: Antisense (179 regions); I: Intergenic (47 regions); H: ncRNA host (27 regions); functional lncRNA: set of 90 regions from 27 lncRNAs with annotated functions in lncRNAdb; codRNA: protein coding transcripts with BLASTN matches (13,034 regions).

### Conserved lncRNAs regions are enriched in translated ORFs

Actively translated sequences show a characteristic three-nucleotide read periodicity in ribosome profiling experiments, allowing the identification of novel translation events (Bazzini et al., 2014; Chew et al., 2013; Ingolia et al., 2009). We used the program RibORF to score read periodicity and uniformity in all ORFs of size 30 nucleotides or longer (Ji et al., 2015). Translated ORFs were defined as those with a RibORF score equal or higher than 0.7, as previously described (Ruiz-Orera et al., 2018) (Figure 4a). The program predicted that 52.05% of all expressed lncRNAs translated at least one ORF, which is in line with previous studies (Calviello et al., 2016; Ji et al., 2015; Ruiz-Orera et al., 2014, 2015). Annotated functional lncRNAs were no exception; we found significant Ribo-Seq signal in 29 out of 30 annotated functional lncRNAs. Except for *TERC*, *Rian*, *Mir124a-1hg*, and *Kcnq1ot1*, the rest of the cases contained small ORFs that appeared to be translated (Additional File 2: Table 3). Virtually all codRNAs with conserved regions were translated; in the case of lncRNAs with conserved regions this fraction was 57.1% (Figure 4b).

**Figure 4.**
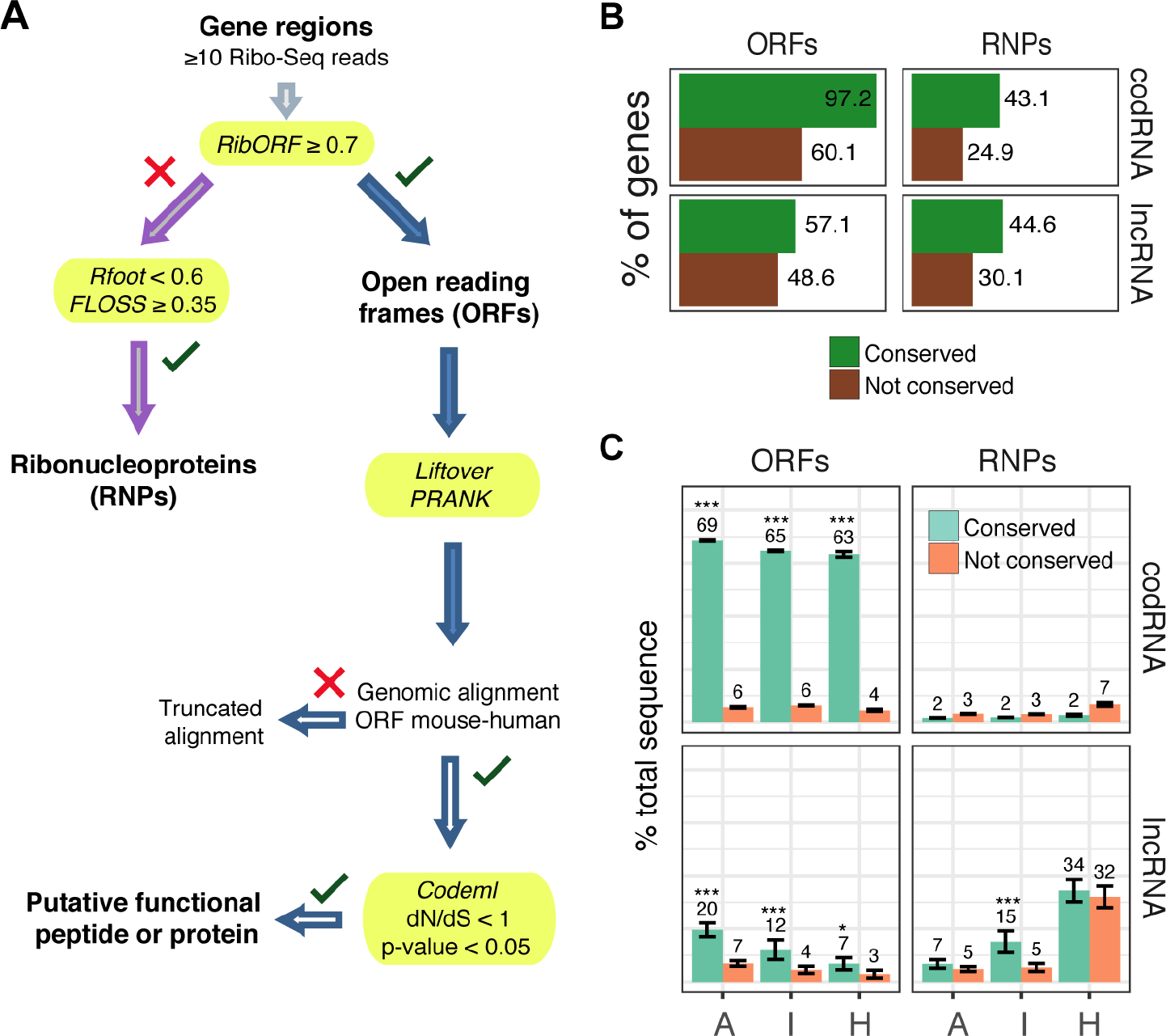
Identification of translated open reading frames and ribonucleoproteins. **A.** Workflow to identify translated open reading frames (ORFs), putative functional proteins, and ribonucleoproteins (RNPs). Ribosome profiling (Ribo-Seq) reads are mapped to candidate gene regions and ORFs with a RibORF score >= 0.7 are defined as translated. Rest of regions with Rfoot uniformity score < 0.6 and FLOSS score >= 0.35 are defined as RNPs. Next, human ORF syntenic regions are extracted with LiftOver and aligned with PRANK, when possible. Truncated alignments are those ones in which less than 50% of the ORF was aligned, or the gap limit is exceeded (33% or 10-nt). Finally, non-truncated alignments are checked for purifying selection signatures with Codeml to identify putative functional peptides or proteins (dN/dS ratio < 1; Chi-square test of dN/dS ratio, p-value < 0.05). **B.** Percentage of conserved and not conserved codRNAs and lncRNAs that contain at least one translated open reading frame (ORFs) or ribonucleprotein (RNPs). Conserved genes show an enrichment in ORFs and RNPs. **C.** Percentage of total sequence that is covered by open reading frames (ORFs), and ribonucleproteins (RNPs), for conserved and non-conserved regions. CodRNA and lncRNA regions showed a significantly higher proportion of ORFs compared to not conserved regions or expected randomly (Test of equal proportions; * p-value < 0.05; *** p-value < 10^−5^). Error bars represent the standard error of the proportion. Categories: A: Antisense; I: Intergenic; H: ncRNA host.

Overall, about 14.1% of the total conserved region in lncRNAs contained ORFs predicted to be translated (122 ORFs), compared to 5.65% for non-conserved regions (370 ORFs). The enrichment of translated ORFs in conserved regions was highly significant (Figure 4c, Test of equal proportions, p-value < 10^−5^). A similar result was observed after discarding regions overlapping other genes in antisense direction (Additional file 1: Figure S3). We also observed that the translated ORFs were more abundant in the 5’ end than in the 3’ end of genes, independently of mouse-human sequence conservation (Additional file 1: Figure S4). This may be related to the ribosome scanning dynamics, starting at the 5’ end of transcripts, and perhaps also to the higher GC content in this part of the gene (Additional file 1: Figure S8), which may favor the presence of ORFs (Vakirlis et al., 2018). The enrichment was consistently observed across the different lncRNA subtypes, with translation occurring more actively in antisense genes than in other lncRNA classes (Figure 4c).

We next investigated if the putative translated ORFs in lncRNA conserved regions showed signatures of selection at the protein level (Figure 4a). Out of the 93 cases in which we could recover and align the corresponding human sequences using genomic alignments, 10 showed a ratio of non-synonymous to synonymous rates (dN/dS) significantly lower than 1 (chi-square test, p-value < 0.05), indicating that this subset of translated products might be functional. The size of the new putative proteins ranged from 19 to 128 amino acids and they were located in uncharacterized lncRNAs (Additional File 2: Table 4). Even though many annotated functional lncRNAs had ORFs in conserved regions, none of them had significant signatures of selection at the protein sequence level. For comparison we also analyzed the signatures of selection in 157 conserved and 38 not conserved codRNA genes encoding small proteins (small CDSs, < 100 amino acids). In this case a much higher proportion of the aligned cases (76 out of 124) showed significant negative selection signatures. These cases included a number of known functional peptides such as Myoregulin (Anderson et al., 2015; Yu et al., 2017), Tunar (Lin et al., 2014), NoBody (D’Lima et al., 2017), or CASIMO1 (Polycarpou-Schwarz et al., 2018), originally annotated as lncRNAs; and other small functional peptides such as Stannin (Buck-Koehntop et al., 2005; Pueyo et al., 2016), or Sarcolipin (Magny et al., 2013; Wawrzynow et al., 1992).

We analyzed PhyloP scores for +1,+2, and +3 positions in codons (see Methods) and we observed that small CDSs had lower conservation values for nucleotides in the third position, which has been generally observed for annotated proteins (Pollard et al., 2010). However, only conserved ORFs with dN/dS-based evidence of negative selection in lncRNAs had the same bias in the third position (Additional file 1: Figure S9). We concluded that, although lncRNA conserved regions are enriched in putatively translated ORFs, probably only a relatively small subset of them are producing functional peptides.

### Identification of RNA-protein interactions

Analysis of fragment length on Ribo-Seq data has revealed differences in the patterns of sequences bound to ribosomes and to small RNAs (Ingolia et al., 2014; Ji et al., 2016). When analyzing the regions covered by Ribo-Seq reads we found that most codRNAs were covered by reads with lengths of 30-32 nucleotides, which correspond to ribosome associations. In lncRNAs the length of the Ribo-Seq reads was more variable, as would be expected if, in addition to translated ORFs, there were non-ribosomal protein-RNA interactions, or ribonucleoproteins (RNPs). The excess of short (< 30 nt) and long (> 32 nt) reads could be clearly observed in intergenic lncRNAs and ncRNA host (Figure 5a). Similar patterns of Ribo-Seq read length were observed in another ribosome profiling experiment from rat when looking at the syntenic regions (Additional file 1: Figure S10).

**Figure 5.**
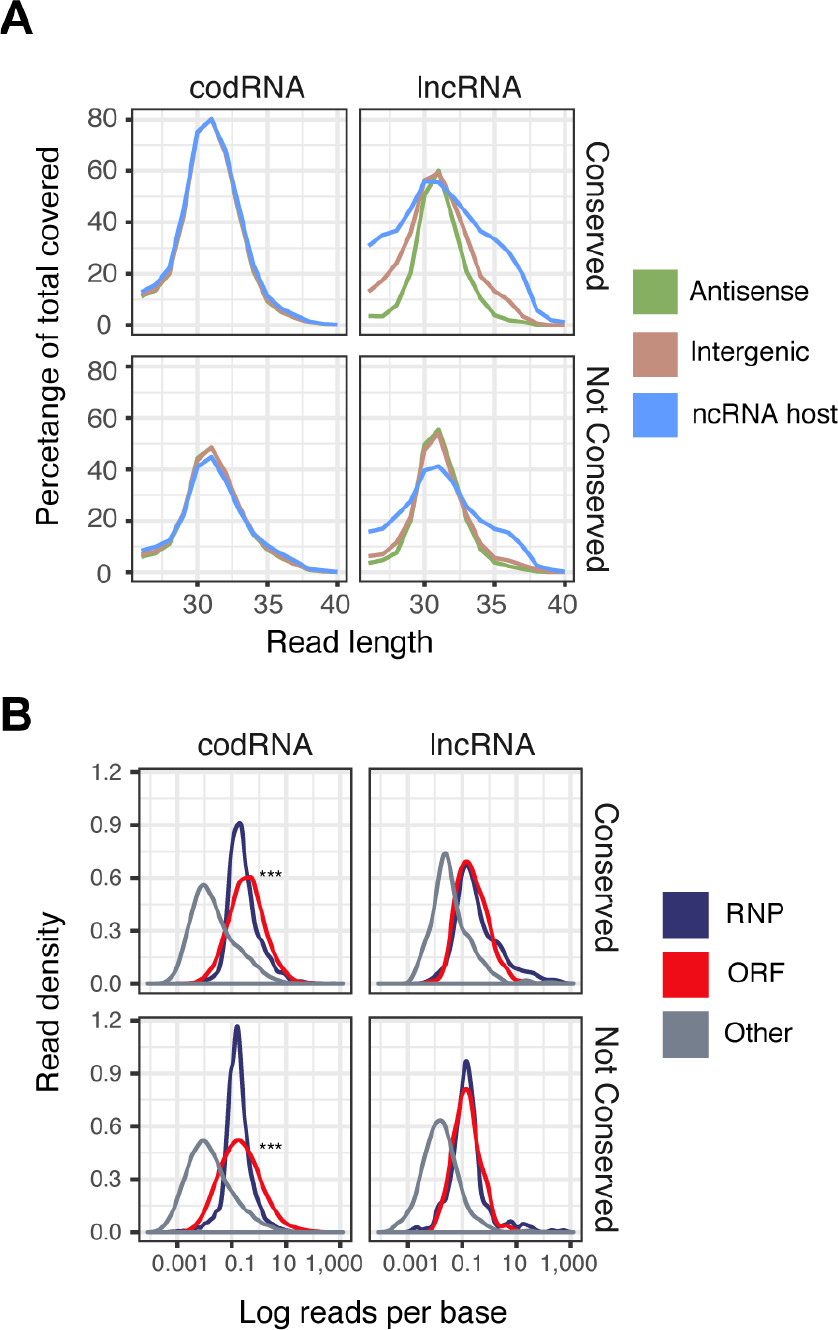
LncRNAs have more heterogenous Ribo-Seq read length. **A.** Fraction of sequence covered by Ribo-Seq that contains reads from a specific length for conserved and not conserved regions in different categories of lncRNAs. While antisense lncRNAs resemble codRNAs in the read distribution, intergenic and ncRNA host regions contain a higher proportion of short and long reads corresponding to non-ribosome associates. **B.** Ribo-Seq read density for regions predicted as ribonucleoproteins (RNP), translated sequences (ORF) and other regions covered by Ribo-Seq. ORFs in codRNAs have a higher read density than the rest of sequences (***. Wilcoxon test, p-value < 10^−5^)

We searched for candidate RNP regions by first identifying regions with low Ribo-Seq read uniformity (< 0.6) with the program Rfoot (Ji et al., 2016), and then checking that the Ribo-Seq reads spanning these regions had lengths which were more compatible with RNPs than with ribosome associations using the previously developed FLOSS methodology (Ingolia et al., 2014). In particular, we selected RNP candidates with a FLOSS divergence score ≥ 0.35 (Figure 4a and Additional file 1: Figure S11). This procedure identified 134 conserved regions in 84 genes that had RNP signatures. This included 21 annotated lncRNAs known to be involved in different functional protein interactions (Additional File 2: Table 3). Among them there were *Malat, Neat1, Meg3*, and *Miat*, known to interact with different protein and splicing factors, and *TERC*, which acts as a scaffold for the telomerase complex. We also found 32 uncharacterized antisense lncRNAs, 12 intergenic lncRNAs, and 19 ncRNA host genes that showed sequence conservation in humans and were associated with RNPs (Additional File 2: Table 2). RNP regions had normalized substitution rates (*k*_*o*_/*k*_*e*_) lower than the simulated sequence evolution control (Wilcoxon test, p-value < 10^−5^), but not different from conserved lncRNA regions in general.

There was an enrichment of RNP signaures in transcripts with at least one conserved region when compared to transcripts with no conserved regions (Figure 4b, Conserved versus Not conserved). The trend was highly significant for intergenic lncRNAs, with 15% of the conserved regions covered by putative RNPs (Figure 4c). Among non-conserved genes with RNP signatures we identified *Firre*, a functional lncRNA that interacts with nuclear factors through a repetitive sequence (Hacisuleyman et al., 2014). In this lncRNA, the predicted RNPs matched the repetitive sequences.

The RNP signatures were clearly lower in codRNAs than in lncRNAs. For example, whereas in lncRNAs read coverage was similar for RNPs and translated ORFs, in codRNAs the translated ORFs had higher coverage (Figure 5b). In conserved lncRNA regions, RNPs and ORFs occupied a similar percentage of the sequence (17.2% and 14.1%, respectively). In contrast, in conserved codRNA regions, RNPs only occupied 1.7% of the sequence, whereas ORFs occupied 65.5% (Additional file 1: Figure S12).

## DISCUSSION

Here we performed an evolutionary analysis of the mouse transcriptome and studied the relationship between evolutionary conservation and the presence of regulatory elements and Ribo-Seq-related features. Several previous studies used regions of predefined genomic synteny to identify homologous regions with primary sequence conservation (He et al., 2015; Hezroni et al., 2015; Li and Yang, 2017; Mohammadin et al., 2015; Ulitsky et al., 2011). These studies showed that lncRNAs were less conserved across distant species than protein-coding genes (Guttman et al., 2009; Marques and Ponting, 2009; Necsulea et al., 2014). However, genomic conservation does not always imply conserved lncRNA expression and/or functionality. LncRNAs are known to have a high expression turnover (Kutter et al., 2012; Neme and Tautz, 2016; Ruiz-Orera et al., 2015) and thus lncRNA expression is often species-specific or limited to very close species, even when genomic syntenic regions can be identified in more distant species (Hezroni et al., 2015). In order to circumvent these limitations here we focused on sequences expressed both in mouse and human, and which had significant sequence similarity by BLASTN, denoting common ancestry. We identified 289 (40.88%) lncRNAs expressed in hippocampus with homology to human transcripts. Conserved regions in lncRNA were usually small; they occupied 8.50% of the total mouse lncRNA sequence length. Although these regions were small, we have to consider that a short region may in some cases be sufficient to carry out the function of the lncRNA (Quinn et al., 2014). In some cases, exon structures located in the 3’ region may be rewired without necessarily affecting lncRNA and/or promoter function, as it occurs with the *Pvt1* gene (Hezroni et al., 2015).

Previous studies found that lncRNAs conserved across different species were more constrained than species-specific lncRNAs (Kutter et al., 2012; Wiberg et al., 2015) or that sequences presumably evolving under no constraints (Marques and Ponting, 2009). It was also reported that putative low-accessibility nucleotides from secondary structure elements showed a depletion of polymorphisms when compared to other exonic and intronic sequences (Pegueroles and Gabaldón, 2016). Here we estimated the nucleotide substitution rate (*k*) from mouse and human lncRNA aligned regions, and compared it to the expected one for sequences evolving under no constraints. We found that, in general, conserved regions in lncRNAs had significantly lower substitution rates than neutrally evolved sequences. This finding is consistent with the existence of evolutionary constraints in lncRNA conserved regions; those positions that are important for the function of the transcript will tend to change less than expected by chance. However, we also have to consider that the mutation rate may be quite heterogeneous in different genomic locations and that this may generate biases that are not related to selection and which are difficult to model.

We found that lncRNA conserved regions were frequently located in the 5’ end of transcripts and that they frequently overlapped with putative promoter sequences. This is in line with previous observations that promoters of conserved mammalian lncRNAs tend to show low sequence divergence (Derrien et al., 2012; Guttman et al., 2009). This pattern may be explained by selection acting to maintain the expression of the gene, but there may also be a certain degree of ascertainment bias, as homology searches will favor the detection of transcripts with conserved promoters even if selection is not acting.

In many cases, regions other than promoters were conserved and associated with low substitution rates. This included 95% of the transcripts hosting small RNAs, which are expected to contain functional RNA molecules, but also 27% of the intergenic and 42% of the antisense lncRNAs. As it has been previously observed that lncRNAs often contained ribosome profilign signatures (Aspden et al., 2014; Bazzini et al., 2014; Guttman et al., 2013; van Heesch et al., 2014; Ingolia et al., 2011; Juntawong et al., 2014; Ruiz-Orera et al., 2014), we hypothesized that evoutionary conservation could be related to the presence of translated ORFs or non-ribosomal ribonucleoprotein particles in the transcripts. We analyzed the patterns of Ribo-Seq in a mouse hippocampus dataset and observed a Ribo-Seq bias towards the 5’ end fraction of both coding and non-coding transcripts. Remarkably, our approach found a very significant enrichment of Ribo-Seq reads in lncRNA conserved regions. The findings were consistent across different expression ranges and species, strengthening our conclusions.

The presence of Ribo-Seq signal in lncRNAs has been previously proposed to be the result of the ribosome scanning of 5’ UTR sequences and the translation of numerous ORFs (Calviello et al., 2016; Ji et al., 2015; Ruiz-Orera et al., 2014), especially in the 5’ end of the RNA (Ingolia et al., 2014). Population analyses on single nucleotide polymorphisms led to the conclusion that many ORFs produce neutral peptides, but some of them are conserved across different species and might translate functional small peptides and proteins (Bazzini et al., 2014; Ruiz-Orera et al., 2018). Small peptides are usually difficult to detect as the small size may hinder the detection by proteomics (Slavoff et al., 2013). As a result, some annotated lncRNAs have only recently been found to translate small functional proteins. This includes cases of lncRNAs previously reported to be functional at the non-coding level, as *Tunar/Megamind* (Lin et al., 2014) or *Mrln* (Anderson et al., 2015; Yu et al., 2017). Here we detected an enrichment of ORFs in conserved regions, which is biased towards the 5’end of the transcript, with antisense lncRNAs showing the highest enrichment. We found at least 10 cases in which the encoded peptide is likely to be functional, and which deserve further investigations. In many other cases the ORFs translated peptides that did not showed signs of functionality, as recently observed for many species- and lineage-specific transcripts (Ruiz-Orera et al., 2018). It is also possible that, in some cases, the ORF may have differed extensively between mouse and human due to the rewire of non-conserved 3’ end exons (Almada et al., 2013). The results are also consistent with the hypothesis that some some translated sequences in lncRNAs might be regulatory ORFs that influence the stability of the transcript (Carlevaro-Fita et al., 2016; Johnstone et al., 2016); in some cases the putative regulatory ORFs may derive from ancient protein-coding genes (Hezroni et al., 2017). In contrast to lncRNAs, most small proteins translated by protein-coding genes showed evidence of selection at the protein level. This included several recently discovered micropeptides, such as Nbdy (D’Lima et al., 2017). As the ribosome profiling data we analyzed was from neural tissues, the newly discovered micropeptides are likely to be enriched in neural functions. The analysis of different tissues and conditions might reveal new functional small peptides that are not expressed or translated in hippocampus. For example, we did not find expression of some recently characterized small functional peptides such as Apela (Pauli et al., 2014), Spaar (Matsumoto et al., 2016), Dworf (Nelson et al., 2016), or Mymx (Zhang et al., 2017), which are expressed in other tissues (Ruiz-Orera et al., 2018).

Although we found many translated ORFs in lncRNAs, many Ribo-Seq reads were distributed along the transcript with low three-frame periodicity and uniformity, two parameters that are used to predict protein translation (Ji et al., 2015). These reads are often the result of non-ribosomal RNA-protein interactions and do not correspond to true 80S footprints. Two different methods have been proposed for the identification of such ribonucleoprotein particles (RNP) signatures: FLOSS, which is based on deviations from the expected RNA length covered by ribosomes (Ingolia et al., 2014) and Rfoot, which selects regions on the basis of low uniformity of the reads (Ji et al., 2016). We reasoned that protein-RNA interactions should display the two types of signatures to be sufficiently reliable, and designed a specific pipeline that integrated the two approaches. The method selected 21 functionally characterized lncRNAs, including intergenic loci as *Malat, Neat1*, or *TERC*, and 19 loci known to host small RNA elements, such as microRNAs or snoRNAs, as well as 44 new unknown candidates. These lncRNAs will be an interesting resource for characterizing novel functional RNA-protein interactions, as they displayed the same level of sequence constraints than functionally characterized lncRNAs. We also found a significant number of RNPs within non-conserved regions; this could be due to promiscuous RNA-protein interactions (Davidovich et al., 2013), the existence of young functional lncRNAs (Durruthy-Durruthy et al., 2015; Heinen et al., 2009; Rigoutsos et al., 2017), or lncRNAs that only contain repetitive, very small, or poorly conserved sequences, as observed for the functionally characterized ncRNA *Firre* (Hacisuleyman et al., 2014) or for some secondary structure elements detected in *Neat1* (Lin et al., 2018).

This study has shown that mouse and human conserved lncRNA sequences show significant evolutionary constraints and a more than two-fold enrichment in ribosome profiling (Ribo-Seq) signatures with respect to non-conserved regions. This includes a number of putative functional micropeptides as well as lncRNAs that contain protein-RNA interaction domains. When we consider translated open reading frames, protein-RNA interaction signatures, putative promoter regions and overlapping antisense exons, our analysis covers 77.4% of the annotated mouse lncRNA sequences with significant homology to human transcripts (Additional file 1: Figure S12). This study integrates disparate data into a common evolutionary framework and builds testable hypotheses about the functions of many lncRNAs.

## EXPERIMENTAL PROCEDURES

### Identification of conserved and non-conserved regions in the mouse transcriptome

We retrieved genome sequences, gene annotations, and regulatory regions (core promoters elements) from Ensembl v.89 for mouse (Flicek et al., 2013). We excluded pseudogenes and sense intronic lncRNAs, as the latter could represent unannotated regions of protein-coding genes. In order to avoid spurious conservation matches due to the presence of repeats and transposable elements, we masked repetitive sequences with RepeatMasker (Smit, AFA, Hubley, R & Green). The masked regions comprised 11.30% of codRNA and 11.56% of lncRNA total sequence. We retained those sequences that had a minimum length of 200 nucleotides and a non-masked sequence length of at least 100 nucleotides or 25% of the total transcript length.

We run BLASTN (Altschul et al., 1997) of the mouse annotated genes against a human transcriptome sequenced at high depth, and comprising both annotated and novel transcripts, from a previous study (Ruiz-Orera et al. 2018). The human transcriptome can be downloaded from http://dx.doi.org/10.6084/m9.figshare.4702375. The BLASTN parameters employed were: -evalue 10^−5^, -strand plus, -max_target_seqs 15, -window_size 12. Next, we defined ‘conserved regions’ in mouse transcripts as the ones showing significant sequence similarity (E-value < 10^−5^) in the human transcriptome. Results were consistent when modifying e-value parameters, as the number of conserved lncRNAs only increased a 4.65% when relaxing the parameter (E-value < 10^−4^) or decreased a 3.36% when making the parameter more stringent (E-value < 10^−6^).

Overlapping BLASTN hits from different transcripts were merged, so every gene had a unique set of conserved non-redundant regions. We defined the gene as codRNA if at least one of the isoforms was protein-coding, otherwise it was defined as lncRNA. We discarded 368 lncRNAs that had homology to sequences annotated as coding in human, as their status was unclear and they might represent unnanotated proteins or pseudogenized lncRNAs. Additionally, if two conserved regions were separated by less than 100 nucleotides we merged them. This was justified by the observation that less than 5% of the annotated coding sequences had internal gaps longer than 100 nucleotides. Using this criterion we were able to recover >95% of total coding sequence length for the cases in which at least one conserved region was found in the translated sequence. This last step had only a minor effect on the median length of the conserved regions in lncRNAs (from 136 nt to 163 nt, Additional file 1: Figure S1). The method identified conserved regions in 19,779 out of 21,416 protein-coding genes (codRNAs) and 1,594 out of 9,734 lncRNAs. Analysis of mouse-human genomic synteny alignments from UCSC (Schwartz et al., 2003) indicated that about 80% of the mouse lncRNA conserved regions could be aligned to human syntenic regions, whereas this fraction decreased to about 50% for nonconserved regions, including many tandem repeats that were masked by BLAST and that are often over-represented in whole-genome alignments (Hezroni et al., 2015).

We quantified the overlap of conserved and non-conserved regions in codRNAs and lncRNAs with regions annotated as promoters in Ensembl, which covered about 1.62% of the genome. These regions are defined by performing peak calling from data corresponding to open chromatin, histone modification, and transcripiton factor binding assays for several cell lines and tissues (Zerbino et al., 2015).

### Null model for sequences evolving under no constraints

In order to test for selection in the aligned mouse and human sequences, we simulated the evolution of sequences along the mouse and human lineages in the absence of selection with Rose (Stoye et al., 1998). As starting sequences we used the annotated mouse lncRNA sequences, as this allowed us to control for sequence composition and GC content. We used the following parameters: HKY model with a TT ratio of 4.26; mouse branch mean substitution 0.34 and indel rate 0.018×2; human branch mean substitution 0.17 and indel rate 0.009×2; indel function: [.50,.18,.10,.08,.06,.04,.04]. Mean substitutions and rate of insertions and deletions values were based on previous estimates (Consortium, 2002; Lunter, 2007; Ogurtsov et al., 2004), using a twofold higher mutation rate in mouse than in human.

After the simulations we run BLASTN, using the same conditions as for real sequences, and recovered the alignments. Up to 59.6% of the mouse simulated sequences had at least one match in the set of human simulated sequences. This corresponded to the 20.8% of the total sequence length.

### Calculation of substitution rates

We estimated the number of substitutions per site (*k* or *k*_*o*_) in BLAST alignments using the maximum likelihood method ‘baseml’ from the PAML package (Yang, 2007) with model 7 (REV). If a position was covered by several BLAST hits we chose the one with the lowest E-value. We discarded *k* values higher than 5, as they might represent computational artifacts. As we observed that *k* values deviate from neutrality in simulations of neutrally evolved sequences with short length, we also computed a normalized substitution rate *k*_*o*_/*k*_*e*_, being k_e_ the expected neutral rate according to the length of the region after modeling a log-linear regression model for short (< 300 nt) and long (>= 300 nt) neutrally evolved sequences separately:

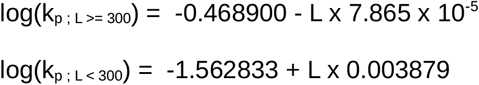

This model was statistically significant for short and long sequences (T-test, p-value < 0.05).

### Classifications of genes based on genomic location or small RNA content

Up to 20% of total sequence length in lncRNAs had exonic overlaps with other genes in the antisense strand. Therefore, we divided conserved and non-conserved regions into ‘overlapping’ and ‘non-overlapping’, depending on whether the region was overlapping with a conserved feature in the other strand (detected by BLAST or annotated as conserved in human in Ensembl Compara). After classifying regions in these 4 different categories, we finally discarded regions shorter than 30 nt and were not considered either as part of the gene, as they might be artifact gaps from homology searches.

Finally, we classified genes in three different categories: ncRNA host, in the case of genes with annotated small RNAs in the exonic structure and/or being annotated microRNA or small RNA host; antisense, in the case of genes having at least one overlapping region, being expressed from bidirectional promoters (closer than 2 kb to an annotated TSS from a antisense protein-coding gene) and/or being annotated as antisense in Ensembl; or intergenic otherwise.

### Analysis of RNA-seq and Ribo-Seq coverage

We used RNA-seq and ribosome profiling data (Ribo-Seq) from mouse hippocampus obtained from Gene Expression Omnibus under accession number GSE72064 (Cho et al., 2015). We merged sequencing replicates to increase the power to detect translated ORFs. We removed reads mapping to annotated rRNAs and tRNAs. Next, we mapped Ribo-Seq (361 million mapped reads) and RNA-seq reads (435 million mapped reads) to the mouse genome (mm10) using Bowtie (v. 0.12.7, parameters -k 1 -m 20 -n 1 --best ‒strata) (Langmead et al., 2009) and we extracted P-sites corresponding to Ribo-Seq reads as done in a previous study (Ruiz-Orera et al., 2018). For comparison, we analyzed Ribo-Seq data from rat brain (rn6, 373 million mapped reads) and human brain (hg19, 50 million mapped reads) obtained from Gene Expression Omnibus under accession numbers GSE66715 (Cho et al., 2015) and GSE51424 (Gonzalez et al., 2014).

Next, we assigned strand-specific mouse reads to the different transcript regions if at least 1bp (RNA-seq) or the computed P-site (Ribo-Seq) spanned the corresponding region. We defined two metrics: a per-base coverage metric based on the number of reads spanning the region per kilobase, and a total coverage based on the percentage of sequence covered by reads.

For genes expressed at very low levels the Ribo-Seq signal may become undetectable. In order to account for this we selected a RNA-seq coverage threshold in which the number of false negatives (annotated coding sequences not covered by RiboSeq reads) was lower than 5% (Additional file 1: Figure S13, RNA-Seq coverage in region ≥ 56.38 reads/kb). In conserved genes, at least one of the conserved regions had to show a coverage above the threshold, or the whole gene was considered as not expressed. Finally, we eliminated 192 lncRNAs located within 4kb from a sense protein-coding gene and/or with evidence of being part of the same gene using RNA-Seq data, these lncRNAs may have been unannotated UTRs.

### ORF translation in conserved and non-conserved regions

We predicted all translated ORFs (ATG to STOP) with a minimum length of 9 amino acids in the transcripts with RibORF (v.0.1) (Ji et al., 2015). Only ORFs with a minimum of 10 mapped Ribo-Seq reads were considered. We used the same score cut-off as in our previosus study (≥ 0.7), which had a reported false positive rate of 3.30-4.16% and a false negative rate of 2.54% (Ruiz-Orera et al., 2018).

Next, we assigned translated ORFs to the different defined regions if at least 10% of the translated sequence spanned a single region. When multiple ORFs spanned one region, we selected the longest one as representative of the region. Consequently, a single ORF could span multiple gene regions, including also discarded ones because of the short length or low expression.

### dN/dS analysis in translated ORFs

We used the UCSC tool liftOver (-minMatch=0.75) (Tyner et al., 2017) to extract the corresponding ORF genomic coordinates in human. For the cases in which we found a matching region, we aligned the ORFs with PRANK (Loytynoja and Goldman, 2005) and we checked how many of these sequences were complete and had the same start-stop codon structure in human, with a resulting coding sequence having at least 50% of the length of the ORF in mouse. Besides, alignment should not contain more than 33% of gaps and aligned length should be longer than 10 amino acids. The remaining ORFs were considered to be truncated.

Next, we used CODEML of the PAML package (Yang, 2007) to compute a dN/dS ratio per complete ORF and we tested whether this ratio was significantly different from 1 by running a fixed omega model. We found 10 mouse and human conserved ORFs in lncRNAs with dN/dS significantly lower than 1, with an adjusted p-value < 0.05.

### PhyloP codon analysis in translated ORFs

We used the GenomicScores package (v. 1.2.2) available at Bioconductor (Puigdevall and Castelo, 2018) to compute the average PhyloP score per codon position (+1, +2, +3) in different sets of translated ORFs. PhyloP is a set of phylogenetic p-values for multiple alignments of 59 vertebrate genomes to the mouse genome. GenomicScores round PhyloP scores using a lossy compresion algorithm. We checked if there was a lower conservation in the third position, as it has been observed in functional proteins (Pollard et al., 2010) due to the degeneracy of the third nucleotide in many codons.

### Analysis of RNA-protein complexes

We used Rfoot (v.0.1) and FLOSS to analyze how many regions in lncRNAs might be involved in RNA-protein complexes (RNPs). Rfoot is a tool that analyzes Ribo-Seq data to identify regions that lack read periodicity and have low read uniformity, and which may correspond to non-ribosomal ribonucleoprotein associates (Ji et al., 2016). FLOSS is a tool that analyzes the distribution of Ribo-Seq read lengths and measures the magnitude of disagreement between distributions to separate ribosome and non-ribosome signals (Ingolia et al., 2014).

We identified putative RNA-protein interaction regions by selecting 60nt windows showing uniformity < 0.6 with a minimum of 10 reads, as done in the original study (Ji et al., 2015). Moreover, we subtracted predicted ORF sequences with a RibORF score ≥ 0.5 and/or read periodicity ≥ 0.66. As ribosome-protected UTR regions could be present in the selected regions, we computed a FLOSS score per region and we defined as RNPs the ones in which the divergence score from ribosome-protected regions was ≥ 0.35. This threshold was selected because only 5% of CDS regions showed a score above 0.35. Subsequently, we merged overlapping regions into a single RNP. This combined approach found RNP associations in 95% of a control set of snRNAs and snoRNAs with 10 or more Ribo-Seq reads, and in only 20% of the 5’ UTRs with the same number of Ribo-Seq reads.

### Evidence of functionality in lncRNAs

We obtained a list of 30 functional mouse lncRNAs expressed in hippocampus by selecting all cases present in lncRNAdb (Quek et al., 2014) and adding four additional known lncRNAs: *Pantr1* (Goff et al., 2015), *Firre* (Hacisuleyman et al., 2014), *TERC* (Feng et al., 1995), and *Norad* (Lee et al., 2016).

### Statistical tests and plots

Plots and statistics was performed with the R package (Team, 2013).

## SUPPLEMENTAL INFORMATION

Additional file 1: File with supplementary information (tables and figures).

Additional file 2: Excel file with properties of the defined lncRNA regions and genes, a list of functionally characterized lncRNAs, and peptide sequences in mouse and human for the 10 functional micropeptides.

Additional file 3: BED file with the coordinates of the lncRNA regions (‘exon’ field), the 492 translated sequences (‘ORF’ field) and the defined RNA-protein interactions (‘RNP’ field).

## ACKNOWLEDGMENTS AND FUNDING

The work was funded by grant BFU2015-65235-P from Ministerio de Economía e Innovación (Spanish Government) co-funded by FEDER (EU), and by grant 2014SGR1121 from Agència de Gestió d’Ajuts Universitaris i de Recerca (AGAUR, Generalitat de Catalunya).

## AUTHOR CONTRIBUTIONS

J.R-O. and M.M.A. conceived the study, interpreted the data and wrote the paper. J.R-O. performed the analyses. M.M.A. coordinated the study. All authors read and approved the final manuscript.

